# Development of a human heart-on-a-chip model using induced pluripotent stem cells, fibroblasts and endothelial cells

**DOI:** 10.1101/2023.12.06.569751

**Authors:** Yun Liu, Rumaisa Kamran, Xiaoxia Han, Mengxue Wang, Qiang Li, Daoyue Lai, Keiji Naruse, Ken Takahashi

## Abstract

In recent years, research on organ-on-a-chip technology has been flourishing, particularly for drug screening and disease model development. Fibroblasts and vascular endothelial cells engage in crosstalk through paracrine signaling and direct cell-cell contact, which is essential for the normal development and function of the heart. Therefore, to faithfully recapitulate cardiac function, it is imperative to incorporate fibroblasts and vascular endothelial cells into a heart-on-a-chip model. Here, we report the development of a human heart-on-a-chip composed of induced pluripotent stem cell (iPSC)-derived cardiomyocytes, fibroblasts, and vascular endothelial cells. Vascular endothelial cells cultured on microfluidic channels responded to the flow of culture medium mimicking blood flow by orienting themselves parallel to the flow direction, akin to *in vivo* vascular alignment in response to blood flow. Furthermore, the flow of culture medium promoted stronger junction formation between vascular endothelial cells, as evidenced by CD31 staining and lower apparent permeability. The triculture condition of iPSC-derived cardiomyocytes, fibroblasts, and vascular endothelial cells resulted in higher expression of the ventricular cardiomyocyte marker IRX4 and increased contractility compared to the biculture condition with iPSC-derived cardiomyocytes and fibroblasts alone. Such triculture-derived cardiac tissues exhibited cardiac responses similar to *in vivo* hearts, including an increase in heart rate upon noradrenaline administration. In summary, we have achieved the development of a heart-on-a-chip composed of cardiomyocytes, fibroblasts, and vascular endothelial cells that mimics *in vivo* cardiac behavior.

## Introduction

Cardiovascular disease poses a significant threat to human health [1], making the creation of disease models related to cardiovascular issues an imperative task. While animal models, such as mice, have traditionally played a pivotal role in this field due to their ease of reproduction and genetic manipulability, their use in cardiovascular research is fraught with two significant inherent problems: inaccuracies in extrapolating data from animals due to differences in cardiac physiology [2, 3], and ethical concerns surrounding animal use. An alternative approach to disease modeling is through cell culture. Typically, cell culture models are two-dimensional (2D), which differs from the three-dimensional (3D) structure of human tissues. Moreover, conventional culture methods struggle to replicate the microenvironment and intercellular communication, including direct contact, paracrine signaling, and extracellular vesicle interactions, which are essential for cellular growth and development [4, 5]. In recent years, advancements in bio-fabrication, microfluidics, and biosensing have introduced 3D culture models [6-9]. Currently, two types of 3D culture models are utilized for constructing cardiac microtissues: organoid and heart-on-a-chip models [7, 8, 10]. In comparison, the heart-on-a-chip model generates cardiomyocytes (CMs) that closely mimic the heart tissue structure [11, 12].

The heart comprises cardiomyocytes, cardiac fibroblasts, and endothelial cells. Cardiac function is not solely dictated by cardiomyocytes but is also influenced by the interactions between cardiac fibroblasts and endothelial cells. Specifically, cardiomyocytes are known to establish gap junctions with fibroblasts, contributing to the establishment of normal electrical excitation in the heart [13, 14]. Additionally, endothelial cells and cardiomyocytes communicate through nitric oxide, fatty acids, and exosomes, which play critical roles in the normal development and postnatal function of the heart [15]. Consequently, co-culturing cardiomyocytes, fibroblasts, and endothelial cells is essential for accurately replicating heart function. Previous research has demonstrated that co-culturing iPSCs with fibroblasts enhances the differentiation of iPSCs into cardiomyocytes [16]. In this study, our objective is to further recapitulate the structure and function of the heart more faithfully by incorporating endothelial cells into this bi-culture system.

In this study, we developed a heart-on-a-chip model by pre-seeding human umbilical vein endothelial cells (HUVEC) in a microfluidic channel. We investigated the effects of medium flow, which mimics blood flow in the vasculature, on endothelial cell morphology and functionality. Endothelial cells exhibited changes in orientation in response to the flow of culture medium, along with the formation of cell-cell junctions mediated by CD31, leading to an enhancement of barrier function. Additionally, by seeding iPSCs and human gingival fibroblasts (HGF) in another microfluidic channel adjacent to the endothelial channel, iPSC-derived cardiomyocytes exhibited increased expression of the ventricular cardiomyocyte marker IRX4 and stronger contractility, compared to iPSC-CM-fibroblast bi-culture. The tri-culture heart-on-a-chip exhibited sensitivity to noradrenaline (NA) and nifedipine similar to that of adult cardiomyocytes. In summary, we developed an iPSC-CM-fibroblast-endothelial tri-culture heart-on-a-chip that better recapitulates heart development and function.

## Results

### Impact of fluid flow on cell alignment

We employed microfluidic chips equipped with two microchannels, namely, top and bottom channels, to create a heart-on-a-chip model. Initially, HUVECs were seeded in the bottom channel of the microfluidic chip. After continuous perfusion of HUVECs for 4 days, the lower ‘vascular’ channel membrane was entirely covered by HUVECs. These HUVECs responded to fluid shear stress by aligning themselves parallel to the direction of medium flow. Under the ‘no flow’ condition, the cells did not exhibit any directional preference (Figures 1A and 1B). In contrast, under the ‘flow’ condition, the cells displayed an elongated, spindle shape with a clear directional preference (Figures 1C and 1D). The results of the directional analysis showed that under the ‘no flow’ condition, HUVECs lacked a specific orientation. However, under the ‘flow’ condition, a distinct directional preference with a peak at 0° was observed. Specifically, the frequency of cell directionality within the range of –20° to 20° was 27.3 ± 3.3% (n = 6) under the ‘no flow’ condition, while it reached 38.6 ± 7.8% (n = 5) under the ‘flow’ condition. (P < 0.01) (Figure 1E). Furthermore, we conducted a directional analysis of cells by varying the flow rates from 0 to 160 μl/h (Figure 1F). As a result, the directional preference toward the direction of fluid flow (0°) became evident at flow rates of 40 μl/h or higher, with a pronounced effect observed at 80 μl/h or above. Additionally, the flow of the medium also had a significant impact on the orientation of F-actin within endothelial cells (Figure 1G). The orientation of F-actin was unclear under the ‘no flow’ condition, while a clear alignment towards the direction of flow was observed under the ‘flow’ condition. Based on these results, a flow rate of 80 μl/h was employed for perfusion hereafter.

**Figure 1.**
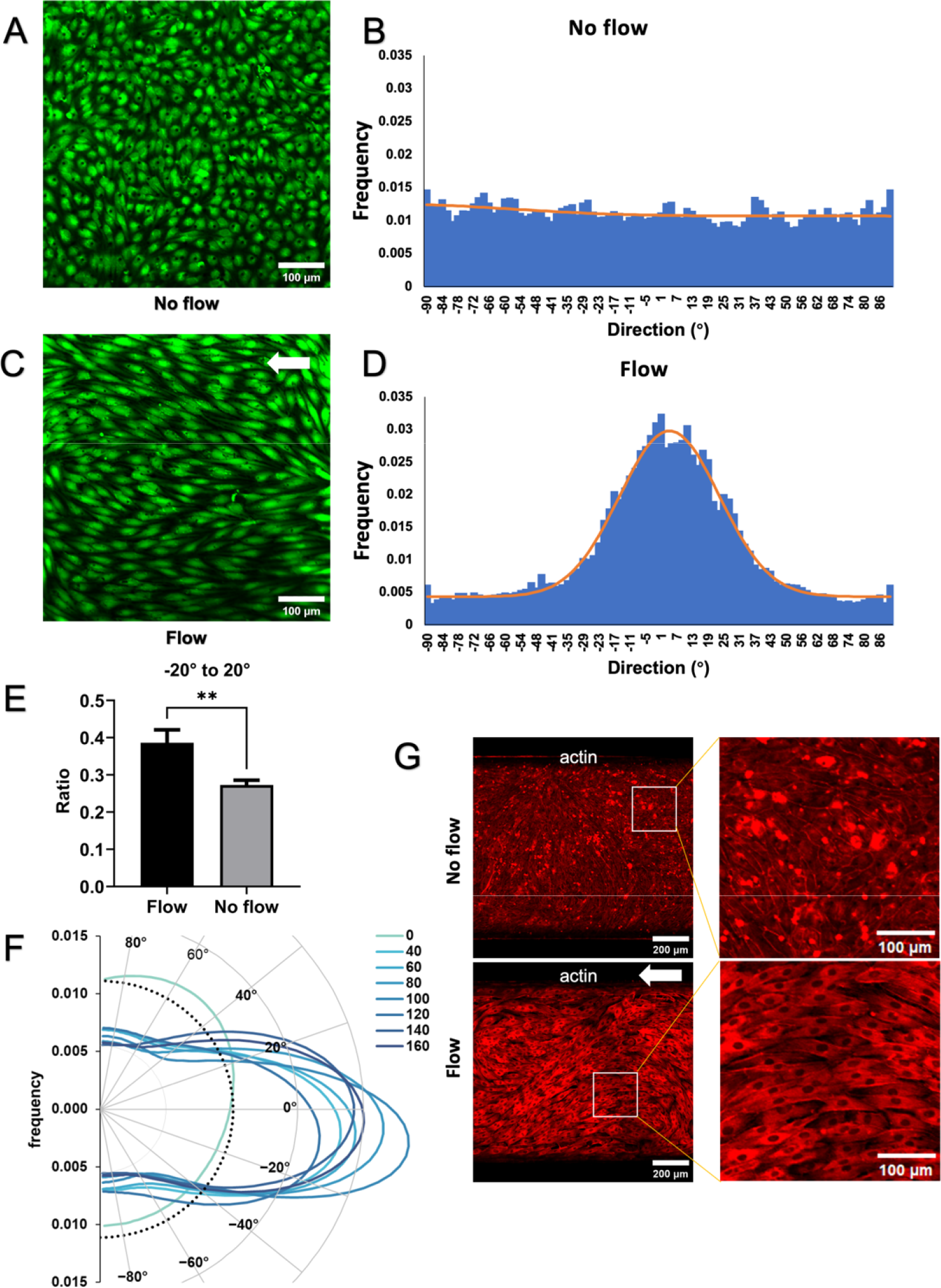
Response of vascular endothelial cells to medium flow. Vascular endothelial cell response to medium flow was examined by seeding HUVECs in microfluidic channels without (A) and with (C) medium flow to study their reaction to fluid shear stress. Frequency histograms of directionality in both groups are depicted in B and D. If there is no orientation in the cell images, the frequency in each bin becomes 0.011. The 0° position signifies the direction of fluid flow in the culture medium. A comparison of total frequencies in the range from −20° to +20° is shown in E. **: P < 0.01. F. Polar representation of directional analysis under varying flow rates of the culture medium is illustrated, with dotted lines indicating a frequency of 0.011 when there is no orientation in the images. G displays a staining image of F-actin in vascular endothelial cells.

### Impact of fluid flow on endothelial cell-cell junction formation

Next, we evaluated the endothelial layer’s barrier function by immunostaining the cell-cell junction protein and quantifying apparent permeability. Immunostaining for the intercellular junction protein CD31 was conducted, revealing an orientation toward the flow direction of endothelial cells similar to that observed in Figure 1 (Figure 2A). Additionally, while CD31 immunofluorescence staining indicated tight junction presence in both groups, the ‘flow’ condition exhibited a significantly higher average fluorescence intensity compared to the ‘no-flow’ condition (127.1 ± 4.8 and 110.2 ± 5.6, respectively, n = 11 for each) (P < 0.05) (Figure 2B).

**Figure 2.**
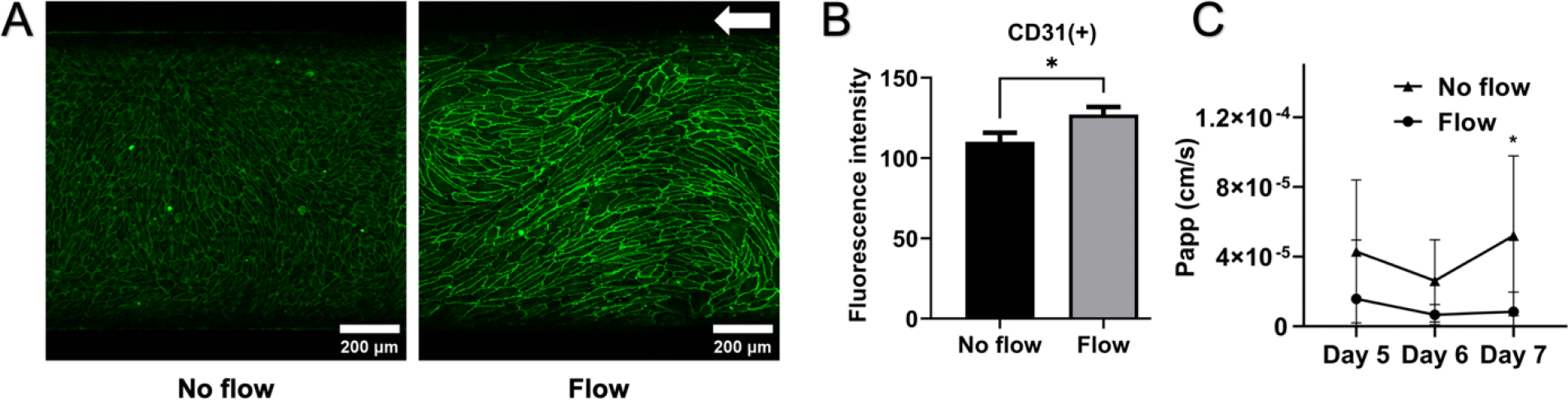
Barrier Function of Vascular Endothelial Cells. A. Comparison of fluorescent immunostaining image of cell-cell junction protein CD31 under flow and no-flow conditions. B. Comparison of fluorescence intensity in CD31 staining. C. Analysis of the apparent permeability (P_app_) of the vascular endothelial cell layer over time. *: P < 0.05.

Furthermore, we determined the apparent permeability from the bottom ‘vascular’ channels to the top ‘cardiac’ channels using Texas Red-conjugated dextran (MW: 3000) (Figure 2C). Between day 5 and day 7 post-seeding of HUVECs, the cells in the flow condition consistently showed a reduction in permeability, decreasing from 1.57 × 10^−5^ ± 3.17 × 10^−5^ cm/s (n = 8, day 5) to 8.32 × 10^−6^ ± 1.07 × 10^−5^ cm/s (n = 10, day 7), representing a 47% reduction. This reduction suggests an enhancement in endothelial barrier function. In contrast, cells in the ‘no flow’ condition exhibited relatively higher permeability of 5.20 × 10^−5^ ± 4.29 × 10^−5^ cm/s (n = 8, day 7) (P < 0.05).

### Endothelial cells enhance the contractility of iPS-CMs

After HUVECs formed a low permeability layer with robust intercellular junctions in the vascular channel, we seeded iPSCs and fibroblasts in the adjacent cardiac channel. Typically, approximately two weeks after initiating differentiation protocol, spontaneously beating CMs were observed in cardiac channels of microfluidic chips. The iPS-CMs continued to exhibit spontaneous contractions until the end of the experimental period, which was the 59th day in this study.

We compared the contractility of iPS-CMs co-cultured with HUVECs (tri-culture group) to those without HUVECs (bi-culture group). Between the 12th and 39th days post-differentiation, we recorded videos of beating CMs in both groups at the same time every 3-4 days (n = 3 biological replicates for each group). From the analysis of 47 recordings (tri-culture group: n = 26, biculture group: n = 21), we observed that the average contractility of iPS-CMs in the tri-culture group was 4.24 ± 1.84, which was significantly higher than that of iPS-CMs in the biculture group (2.87 ± 2.08) (P < 0.05) (Figures 3A and 3B).

**Figure 3.**
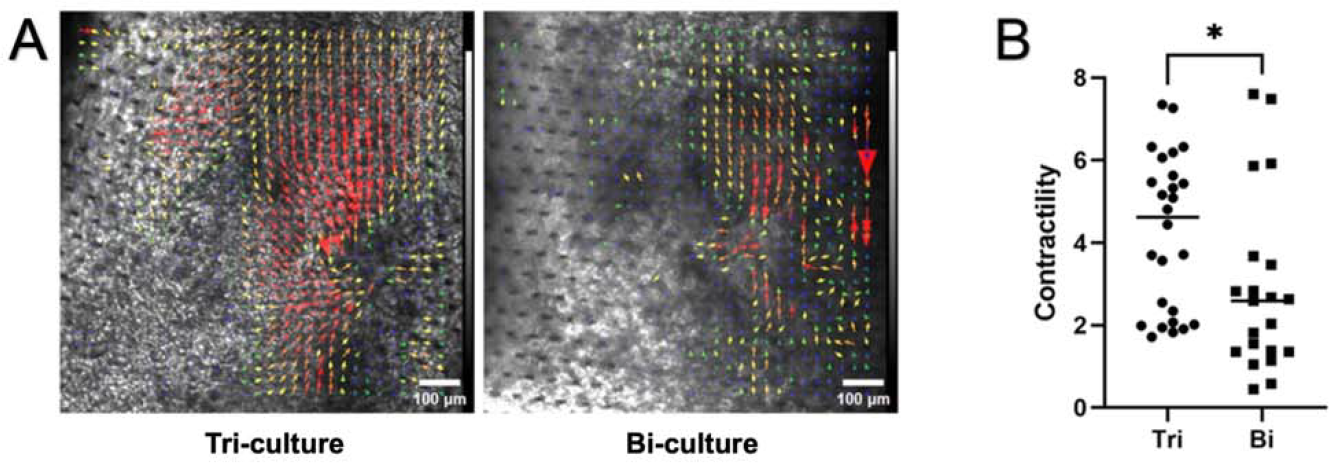
Analysis of cardiac tissue contractility A. Comparison of the vector field of cardiac contractility between tri-culture and bi-culture conditions. Longer vectors indicate higher contractility at the respective sites. B. Comparison of the sum of cardiac contractility between tri-culture and bi-culture conditions. *: P < 0.05.

### Endothelial cells promote cardiac marker expression in iPS-CMs

To investigate whether the high contractility in the tri-culture group is supported by the expression of cardiomyocyte markers, flow cytometry analysis of cardiomyocyte marker expression was performed. In the negative control (HUVEC monoculture), the expression rate of the cardiomyocyte marker cardiac troponin t (cTnT) was 0.1%, whereas it reached 64.8% in a biculture sample and 92.8% in a tri-culture sample (Figure 4A). Although cTnT expression showed a trend of being higher in the tri-culture group (64.6 ± 22.9%, n = 5) as opposed to the bi-culture group (55.9 ± 6.6%, n = 6), this difference did not reach statistical significance (Figure 4B). To explore the localization of cTnT expression in the cardiac tissue, we conducted immunostaining. The tissue in the cardiac channel is relatively thick, around 0.2 mm, and conventional immunocytochemistry protocols didn’t produce satisfactory staining images. Consequently, we prepared paraffin-embedded sections of cardiac tissue for immunostaining. This revealed cTnT localization on sarcomeres, showing a high degree of alignment similar to the *in vivo* heart (Figure 4C).

**Figure 4.**
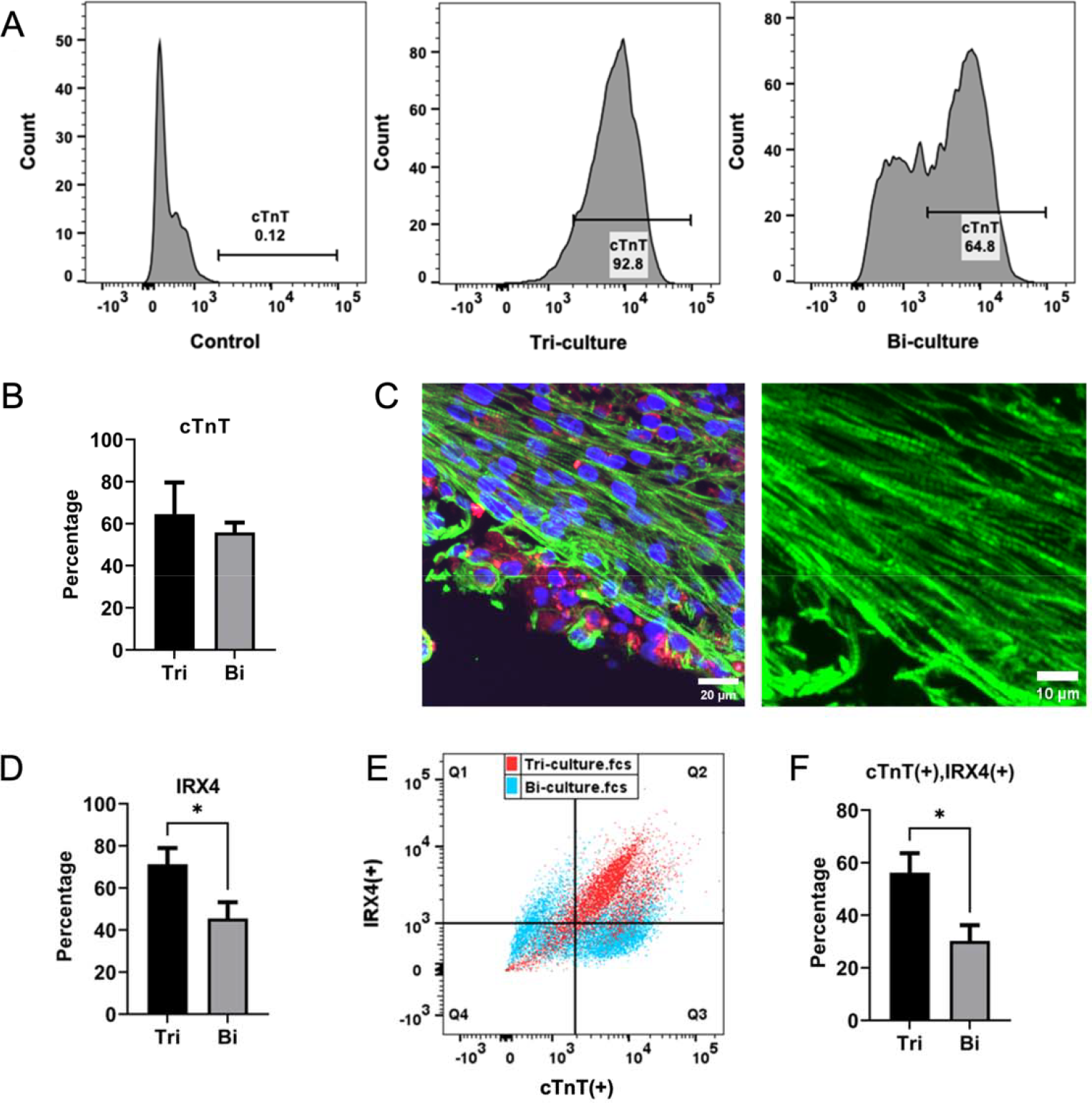
Expression analysis of cardiac marker proteins A. Flow cytometry analysis of cTnT expression in cardiac marker proteins. Control: non-differentiated iPSCs. B. Comparison of cTnT expression levels between the tri-culture and biculture groups. C. Immunostaining image of a paraffin section of cardiac tissue. Green: cTnT, Red: F-actin. In the enlarged image (right), distinct sarcomere structures with clear orientation are observed. D. Comparison of the expression levels of ventricular cardiomyocyte marker IRX4 in the tri-culture and bi-culture groups. E. Analysis of cTnT and IRX4 expression levels by flow cytometry. Red: tri-culture, Blue: bi-culture. F. Comparison of the number of cells co-expressing cTnT and IRX4 between the tri-culture and bi-culture groups. *: P < 0.05.

Following that, we conducted an expression analysis of the ventricular cardiomyocyte marker IRX4. The tri-culture group exhibited significantly higher IRX4 expression compared to the biculture group (P < 0.05, as shown in Figure 4D). Additionally, the percentage of cells coexpressing cTnT and IRX4 was notably elevated in the tri-culture group (56.3 ± 14.7%, n = 5) in contrast to the bi-culture group (30.2 ± 13.5%, n = 6) (P < 0.05) (depicted in Figures 4E and 4F).

### Response of iPS-CMs to noradrenaline

To observe the physiological response of cardiac channel tissues, we administered NA, a β-adrenergic agonist (Figure 5). We observed that NA increased the heart rate of iPSC-CMs in a dose-dependent manner (Figure 5B). The dose-response relationship of NA was fitted with a sigmoid curve allowing a variable slope, resulting in a Hill coefficient of 0.56 and an EC_50_ of 24.8 μM (95% confidence interval: 0.73 - 83.88 μM, n = 4). When the NA concentration was elevated to 1 mM, it resulted in a heart rate of 139.5 ± 10.4, which was 2.61 times the baseline. While NA slightly increased the contractility of cardiac tissue, this effect was modest compared to the increase in heart rate (Figure 5C).

**Figure 5.**
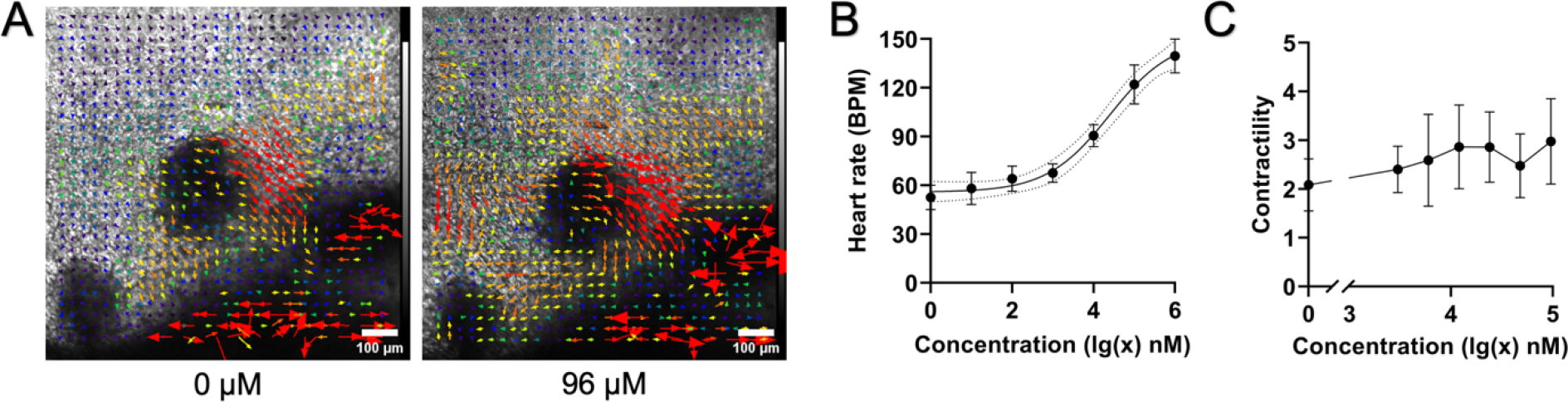
Response of cardiac tissue to noradrenaline (NA) A. Vector field analysis of cardiac tissue contractility before and after NA administration. B. Dose-response curve of NA on heart rate. Estimated EC_50_ is 24.8 μM based on sigmoid curve fitting. C. Dose-response curve of NA on contractility.

### Response of iPS-CMs to nifedipine

Subsequently, we administered nifedipine, a Ca_v_1.2 channel inhibitor, to cardiac channel tissue (Figure 6). Nifedipine, ranging from 0 nM to 1000 nM, did not significantly impact heart rate; however, it elicited a dose-dependent and notably significant decrease in contractility (Figures 6A–6C).

**Figure 6.**
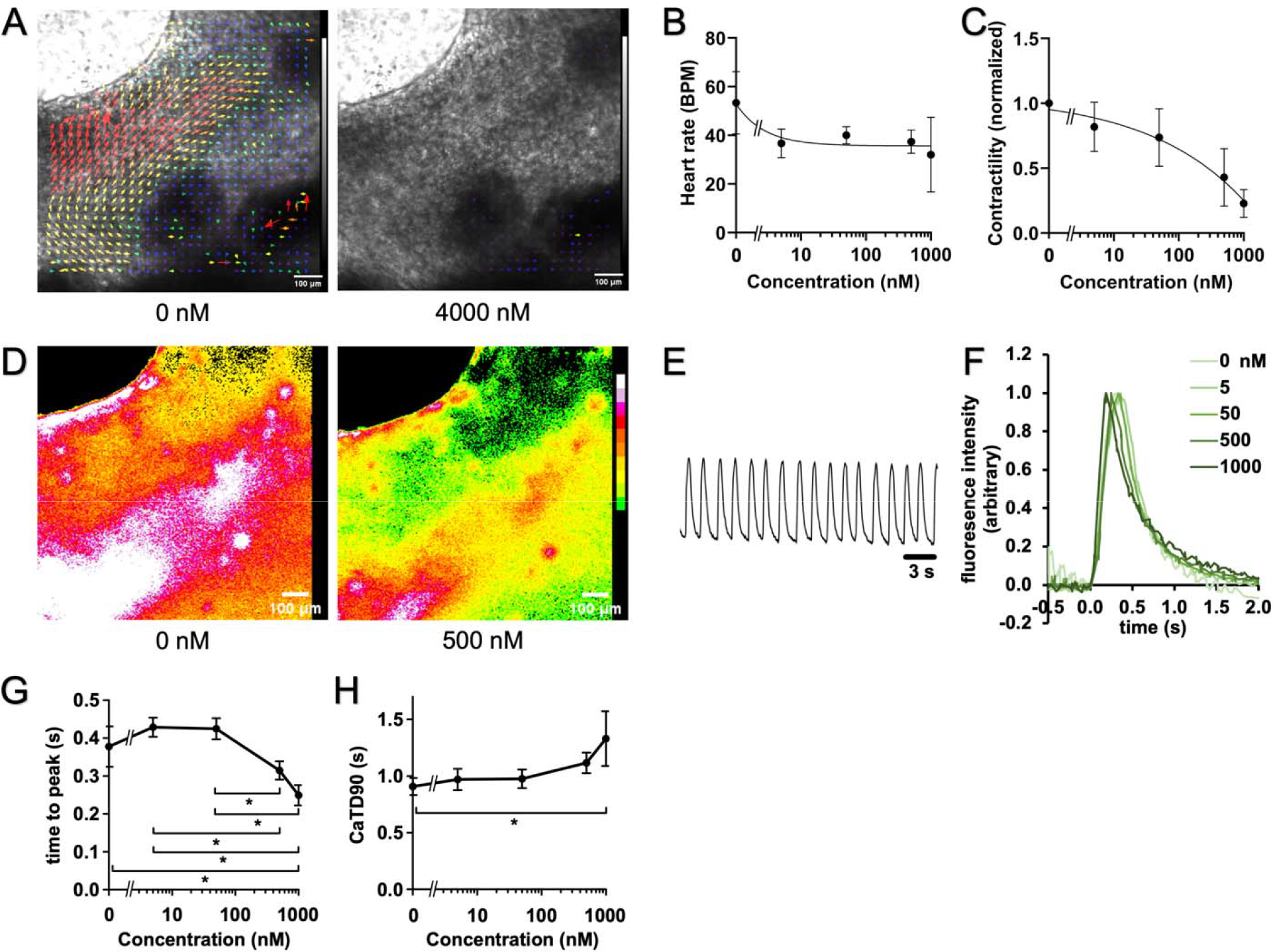
Response of cardiac tissue to nifedipine A. Vector field analysis of cardiac tissue contractility before and after nifedipine administration. B. Dose-response curve of nifedipine on heart rate. C. Dose-response curve of nifedipine on contractility. D. Intracellular calcium ion levels in cardiac tissue before and after nifedipine administration. E. Calcium transient (CaT) trace of cardiac tissue without nifedipine administration. F. Overlay of CaT curves for nifedipine administration at 0, 5, 50, 500, and 1,000 nM. Fluorescence intensity is normalized at the level of peak values. G. Dose-response curve of nifedipine for time to peak in CaT curves. H. Dose-response curve of nifedipine for CaT duration at 90% recovery (CaTD90). *: P < 0.05.

Moreover, we examined intracellular calcium ion dynamics in cardiac tissue using the calcium-sensitive dye Cal-520. iPS-CMs displayed a periodic, coordinated pattern of calcium influx synchronized with their contraction under a fluorescence microscope (Figures 6D and 6E). Fluorescence imaging of calcium transients revealed a consistent pattern of intracellular fluorescence level changes correlated with myocardial contraction. Specifically, fluorescence intensity increased during systole, followed by a decrease during diastole, ultimately returning to baseline. The interval between transient peaks remained consistent throughout the measurement duration (approximately 30 s), with no observed arrhythmias.

Upon normalizing the peak height of the calcium transient, the time to peak decreased in a dosedependent manner with nifedipine (Figures 6F and 6G). In contrast, CaTD90, representing the calcium transient duration at 90% recovery, increased in a dose-dependent manner (Figures 6F and 6H).

## Discussion

Presently, tissue engineering strategies involve 3D reconstruction and incorporate organoids or micro-bioreactors as complementary elements to conventional cell culture models. Moreover, utilizing iPSCs for CM differentiation not only addresses ethical concerns linked to animal models but also facilitates the sourcing of cells from both healthy donors and patients with specific conditions, allowing for a more faithful reproduction of the original donor’s genotype [17, 18]. Additionally, 3D culture models more accurately mimic the normal physiological and anatomical structures of the human body [19], rendering them more representative of organ system physiology.

In this study, we have developed a 3D heart-on-a-chip model using iPSCs, fibroblasts, and endothelial cells, designed to mimic the anatomical structure of cardiac tissue. As demonstrated in previous studies, including our own [16, 20], co-culture with fibroblasts enhances the contractile function of the heart. The addition of vascular endothelial cells to these bi-culture systems in this study holds significant importance, as it brings the heart model even closer to the structure and function of the *in vivo* heart.

In our bodies, vascular endothelial cells are constantly exposed to blood shear stress, initiating intracellular signaling pathways that induce changes in cellular behaviors and functions closely associated with endothelial cell functionality [21]. We applied shear stress to endothelial cells through perfusion using a peristaltic pump. Under shear stress stimulation, endothelial cells aligned parallel to the direction of the flow. The orientation of these endothelial cells is known to occur through the remodeling of F-actin, which constitutes stress fibers [22]. In this study, a clear orientation of flow-induced F-actin was observed.

Vascular permeability is closely related to the pathophysiology of atherosclerosis and is crucial for the manifestation of normal vascular function. At the branching points of blood vessels with a high incidence of atherosclerotic plaque formation, it is known that the permeability of large molecules such as low-density lipoprotein is elevated [22]. In this study, a decrease in apparent permeability was observed under shear stress induced by perfusion of the culture medium. The reduction in vascular permeability is believed to be, in part, attributed to the increased expression of CD31. It is well-documented that shear stress induced by flow upregulates the expression of CD31 [23, 24], a cell-cell junction protein in vascular endothelial cells, and this was observed as an increase in immunostaining intensity in this study. This suggests that the heart-on-a-chip model developed in this study faithfully recapitulates *in vivo* vascular function.

Cardiomyocytes derived from iPSCs often lack stable spontaneous contractions in monoculture conditions [25]. However, in contrast, the cardiac tissue generated through tri-culture in this study demonstrated remarkably stable spontaneous contractions. Additionally, co-culturing with endothelial cells led to increased contractility in iPS-CMs. This heightened contractility coincided with an upregulation in the expression of the ventricular cardiomyocyte marker protein IRX4. Hence, it is suggested that the improved contractility of iPS-CMs through co-culture with vascular endothelial cells likely results from the promotion of differentiation from iPSCs to ventricular cardiomyocytes.

One of the crucial features of cardiomyocytes is their responsiveness to neurohormonal stimulation, such as that induced by norepinephrine [26]. The cardiac tissue in this study demonstrated a dose-dependent positive chronotropic effect in response to noradrenaline. Additionally, another characteristic of cardiomyocytes is a reduction in contractility with the inhibition of calcium channels. In the human heart, inhibiting dihydropyridine calcium channels is known to weaken the contractility of ventricular muscle [27, 28]. Consistently, in this study, the administration of nifedipine, a dihydropyridine calcium channel blocker, led to a reduction in the contractility of cardiac tissue. These findings suggest that the cardiac tissue in this study closely mirrors the functionality of the human heart.

Furthermore, a more detailed analysis of calcium dynamics revealed changes in the calcium transient waveform following nifedipine administration. Specifically, the time from the onset to the peak of the calcium transient was shortened in a nifedipine concentration-dependent manner, while CaTD90 was prolonged. The duration of the calcium transient correlates with the duration of the action potential, and these are associated with the QT interval on an electrocardiogram [29]. Therefore, it is expected that the heart-on-a-chip developed in this study can contribute to investigating human cardiac toxicity, such as the QT-prolonging effects of drugs.

Overall, we have developed a heart-on-chip model mirroring the structure of myocardial tissue, incorporating cardiomyocytes, fibroblasts, and vascular endothelial cells through the utilization of microfluidic chip and 3D culture technology. This heart-on-a-chip exhibits key physiological features of the heart, such as alterations in permeability to blood flow by vascular endothelial cells, spontaneous contractions of cardiac tissue without pacing, and responsiveness to β-adrenergic agonists and L-type calcium channel inhibitors. Moreover, it presents notable advantages, allowing real-time monitoring of cardiac contraction and calcium ion dynamics, facilitating in-depth analyses of contraction vector fields and calcium transient waveforms. The potential applications of this heart-on-a-chip encompass human drug efficacy and toxicity testing, as well as contributing to personalized medicine through the use of patient-derived iPSCs.

## Materials and methods

### Cell maintenance culture

The protocols for maintaining human iPSCs and HGFs are detailed in [16]. iPSCs (201B7 cells derived from dermal fibroblasts) were cultured in StemFit AK02N medium (Cat. #: RCAK02N, Ajinomoto, Japan). Experiments were conducted using iPSCs within the 3rd and 15th passages. For HGF maintenance culture, low-glucose Dulbecco’s Modified Eagle Medium (Cat. #: 11885-084, Thermo Fisher Scientific, MA, USA) supplemented with 10% fetal bovine serum and 1-mol/l-HEPES Buffer Solution (Nacalai tesque, Kyoto, Japan) was employed. All experiments were conducted using HGFs from the 11th to the 20th passages. HUVECs (Cat. #: C2519A, Lonza, Basel, Switzerland) were cultured in Endothelial Cell Growth (ECG) Medium supplemented with 2% v/v fetal calf serum (Cat. #: C-22111, PromoCell, Germany). Subculturing and maintenance cultures were performed using 3 × 10^5^ cells at 37°C in a 5% CO_2_ atmosphere. For experiments, only HUVECs within the passage numbers of 3 to 8 were utilized.

### Cell culture on microfluidic chips

For microfluidic chip experiments, we utilized the Emulate Chip S1 (Chip S1, Emulate, Boston, USA), made of polydimethylsiloxane. This chip comprises a rectangular upper channel measuring 1 mm in width, 1 mm in height, and 17 mm in length, along with a rectangular lower channel measuring 1 mm in width, 0.2 mm in height, and 17 mm in length. The upper and lower channels are separated by a thin film with a thickness of 50 μm, featuring a honeycomb-like arrangement of pores with a diameter of 7 μm, spaced at 40 μm intervals.

Before cell seeding, the chip surface was coated with growth factor-reduced Matrigel (Cat. #: 354230, Corning, USA) to enhance cell adhesion. Initially, the top channel was filled with ECG medium. Subsequently, 200,000 HUVECs in 30 μl ECG medium were pipetted into the bottom channel. The chips were then flipped and incubated overnight in an incubator. The following day, the chips were flipped back and connected to perfusion tubings (Cat. #: SC0188, Ismatec, USA), inlet medium reservoirs, effluent collection containers, and a peristaltic pump (IPC-N 24, Ismatec, USA). The inlet reservoirs were filled with ECG medium. For microfluidic chip culture, Penicillin-Streptomycin Solution (Cat. #: 168-23191, Wako, Japan) and Amphotericin B (Cat. #: A2942, Sigma, USA) were added to the culture medium at a 1:100 dilution each. The perfusion rate was set to 80 μl/h unless otherwise specified.

On the 8th day, a mixture of 100,000 iPSCs and 17,000 HGFs was seeded into the top channel (Figure 7). Perfusion was paused overnight to facilitate cell adhesion. The next day, the culture medium in the top channel inlet reservoir was replaced with Essential 8 medium (Cat. #: A1517001, ThermoFisher, USA), and perfusion commenced. Differentiation of iPSCs into cardiomyocytes was induced using the PSC Cardiomyocyte Differentiation Kit (Cat. #: A2921201, ThermoFisher, USA). Three days after starting perfusion with Essential 8 medium, the culture medium in the top channel inlet reservoir was changed to the differentiation medium A (component of the kit), and perfusion was initiated. After 48 hours, the culture medium in the top channel inlet reservoir was switched to the differentiation medium B, and perfusion was performed. Another 48 hours later, the culture medium in the top channel inlet reservoir was changed to the cardiomyocyte maintenance medium, and continuous perfusion was maintained thereafter.

**Figure 7.**
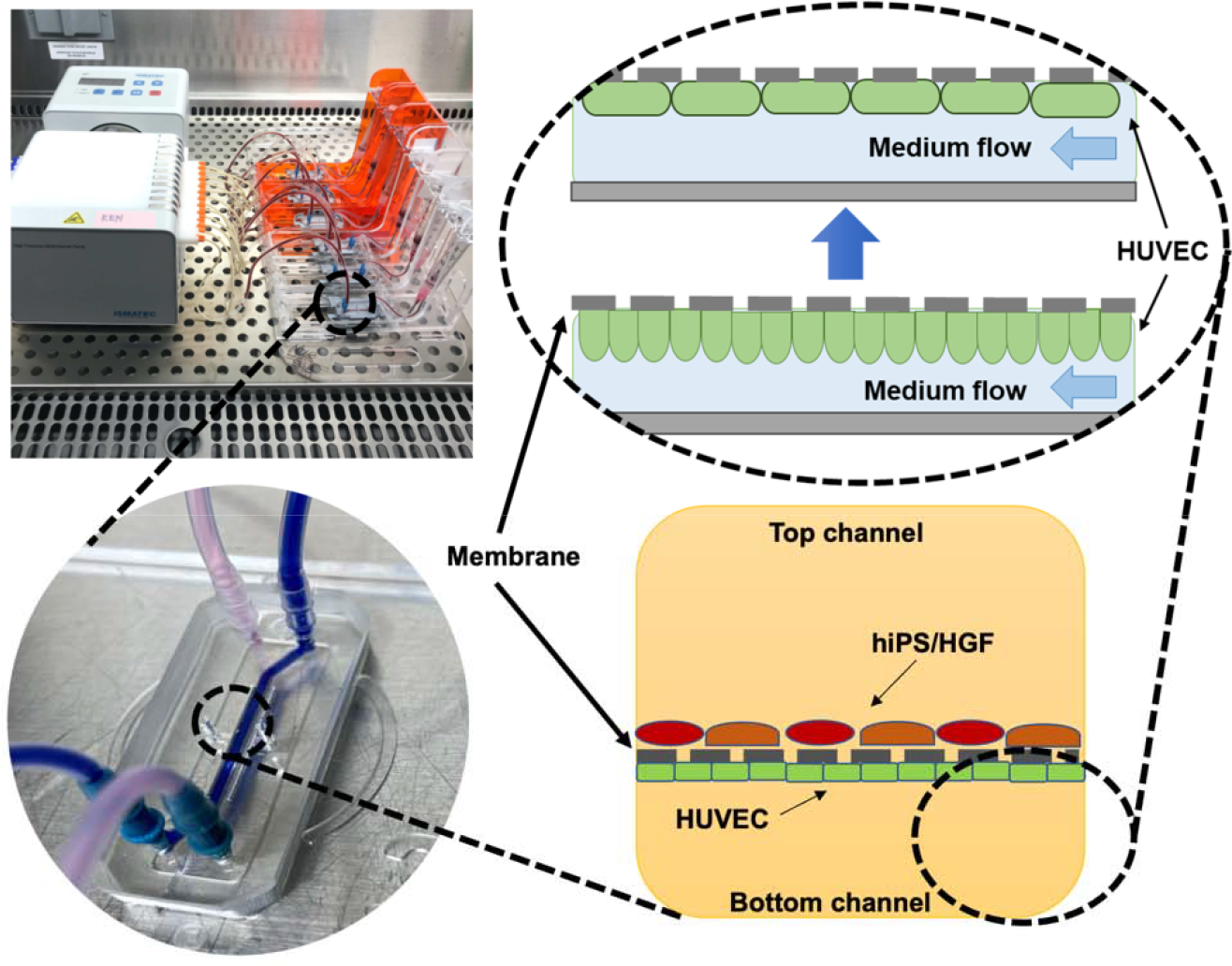
Perfusion system and internal structure of microfluidic chip.

### Permeability Assay

The barrier function of endothelial cells was assessed using the method outlined by Pediaditakis et al. [30]. Texas Red-conjugated dextran (MW: 3000) (Cat. #: D3329, ThermoFisher, USA) served as a fluorescent tracer. The culture medium in the inlet reservoir of the bottom channel was replaced with a medium containing 5 μg/ml dextran and perfused for 2 hours. The effluent solutions collected from the top and bottom channels were used for fluorescence measurements using a plate reader (Flexstation 3, Molecular Devices, USA). Subsequently, the apparent permeability of the dextran was calculated using the formula described in [30].

### Contractility analysis of cardiac tissue

The contractility analysis of cardiac tissue is detailed in a separate publication [16]. In summary, video recordings of beating cardiac tissue were captured at a rate of 20 frames per second using a phase-contrast microscope (BZ-X710, KEYENCE, Japan) at room temperature. A discrete two-dimensional vector field of displacement, with a 16 pixels by 16 pixels grid, was analyzed at each frame. The maximum size of the displacement vector over frames was determined for each grid, and the sum of these maximum sizes across all grids was defined as the contractility of the cardiac tissue.

### Calcium imaging

The calcium indicator Cal-520 (AAT Bioquest, USA) was introduced into the culture medium at a final concentration of 5 μM and perfused for 1 hour, exposing the cells to the compound. Following this, fluorescence images at a wavelength of 520 nm were captured using a fluorescence microscope (ECLIPSE TE2000-U, Nikon, Japan).

Directionality analysis utilized the Cal-520 application mentioned above to enhance the visualization of endothelial cells. Power spectrum analysis was conducted by utilizing the Directionality plugin in ImageJ [31] to Fourier-transform the acquired calcium imaging images.

In the analysis of calcium dynamics in cardiac tissue, fluorescence video imaging was conducted for 30 seconds at an interval of 134 ms at room temperature. Subsequently, to enhance the temporal resolution of the calcium transient (CaT) waveforms, 10 CaT waveforms recorded over 30 seconds were overlaid (Figure 8). Specifically, the first CaT waveform served as a template, and the second CaT waveform was superimposed to align the peak positions. Next, the second CaT waveform was shifted to minimize the difference in fluorescence intensity values at each time point relative to the first CaT waveform. Using a similar approach, overlaying 10 CaT waveforms yielded a high-temporal-resolution CaT curve.

**Figure 8.**
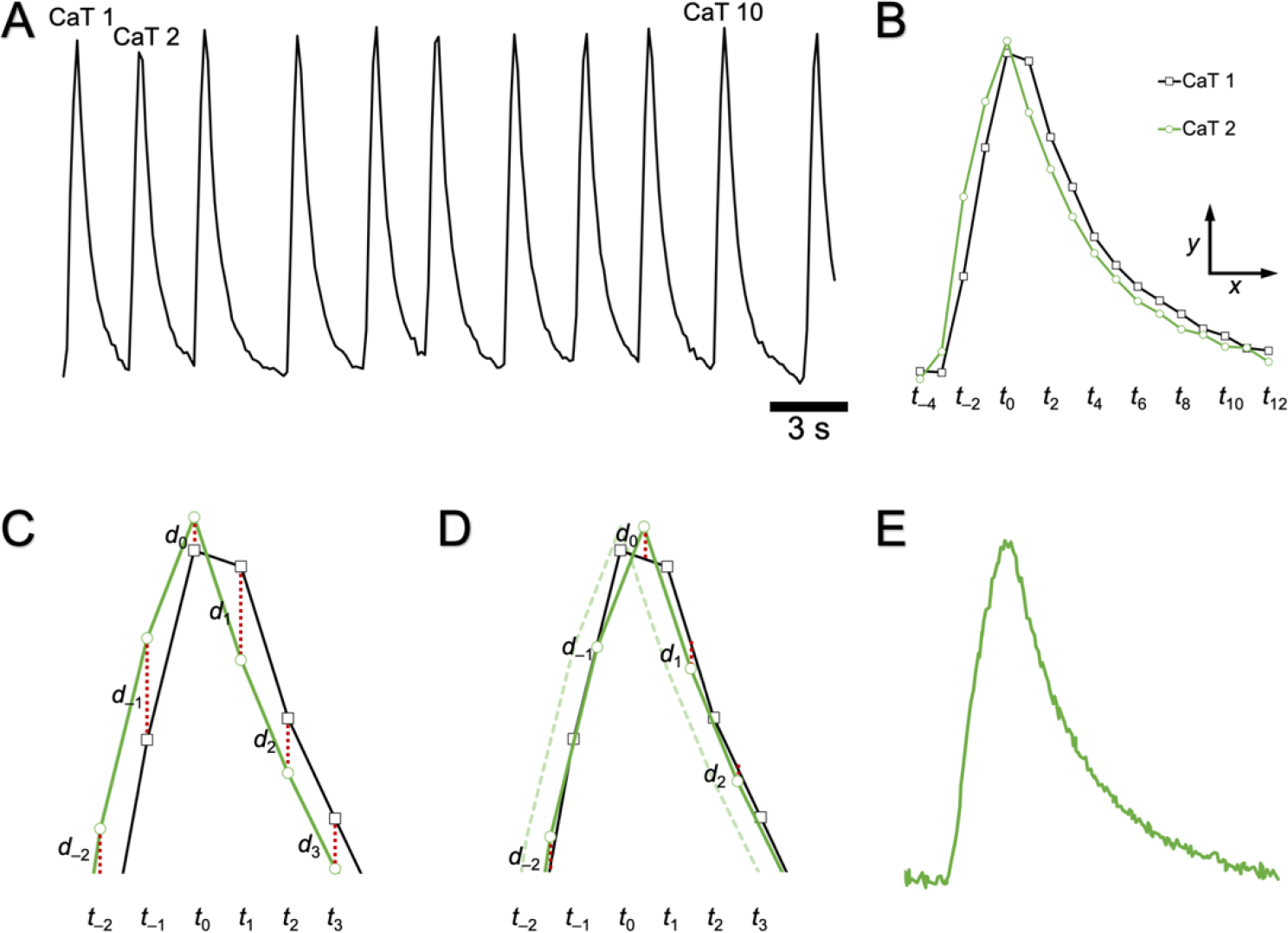
Method to increase temporal resolution of CaT curves A. Typical CaT trace recorded at an imaging interval of 134 ms. B. Overlay of the first (CaT 1) and the second (CaT 2) CaT curves from the trace in A. *t*_*n*_ indicates the timeframe of imaging. C. Magnified view of B. *d*_*n*_ represents the distance between CaT 1 and CaT 2 curves at each *t*_*n*_. D. Diagram showing the movement of CaT 2 in the *x* and *y* directions to minimize the sum of *d*_*n*_. E. CaT curves obtained by overlaying 10 curves. In principle, the temporal resolution is increased by 10 times.

### Staining of CD31 and F-actin

For immunofluorescence staining, cells underwent perfusion with 500 μl of cold Dulbecco’s phosphate-buffered saline (DPBS), followed by fixation with 500 μl of 4% paraformaldehyde (PFA) and permeabilization with 500 μl of 0.125% Triton X-100 (Nacalai tesque, Kyoto, Japan) at a constant rate of 2,000 μl/h using a peristaltic pump. Subsequently, 1.5% bovine serum albumin (BSA; Sigma Aldrich, A9418) in DPBS was perfused for blocking for 1 hour at a rate of 80 μl/h. Following this, the microfluidic chips were disconnected from the perfusion system. A staining solution consisting of rabbit anti-CD31 primary antibody (Cat. #: ab28364, Abcam, UK) diluted 1:100 in the blocking buffer (1.5% BSA in DPBS) was applied to the bottom channel of the chip. The chip was then refrigerated at 4°C overnight. After washing the cells with 200 μl/channel DPBS three times, a staining solution containing goat anti-rabbit secondary antibody conjugated with Alexa Fluor 488 (1:1000 dilution; Cat. #: A-11008, Invitrogen, USA) and ActinRed 555 ReadyProbes Reagent (1:50 dilution; Cat. #: R37112, Invitrogen, USA) was applied to the bottom channel and incubated at room temperature for 1 hour. Fluorescence images were captured using a confocal microscope (FLUOVIEW FV1000, OLYMPUS, Japan).

### Flow cytometry

For flow cytometry analysis of cardiac tissue, the cells were dissociated into single cells by treating them with 180 μl of 0.25% Trypsin/DMEM (1:2) multiple times for 10 minutes, and then collected in a 15 ml centrifuge tube. After centrifugation at 300 × g for 5 minutes, the supernatant was aspirated, and the cells were fixed with a 4% PFA solution at room temperature for 20 minutes. The cells were then centrifuged again, and the resulting cell pellet was resuspended in 200 μl of ice-cold DPBS. Subsequently, the cells were permeabilized with 0.125% Triton X-100 for 20 minutes and blocked with 1.5% BSA in PBS at 4°C for at least 1 hour.

After collecting the cells by centrifugation, they were incubated with a 90 μl primary antibody solution containing mouse anti-cTnT primary andibody (1:100; Cat. #: MA5-12960, Invitrogen, USA) and rabbit anti-IRX4 primary antibody (1:100; Cat. #: PA5-97879, Invitrogen, USA) in the blocking solution at 4°C overnight. Following DPBS washing, the cells were incubated with a secondary antibody solution containing goat anti-mouse secondary antibody conjugated with Alexa Fluor 488 (1:1,000; Cat. #: A32723, Invitrogen, USA) and donkey anti-rabbit secondary antibody conjugated with Alexa Fluor 647 (1:1000; Cat. #: ab150075, Abcam, UK) in the blocking solution at room temperature for 1 hour. Finally, the cells were analyzed using the FACSAria3 cell sorter (BD Biosciences, USA), with 10,000 events recorded for each sample.

### Paraffin section staining

To fix the cells, 200 μl of 4% PFA was applied to the top channel of the microfluidic chip and incubated at 4°C for 12 hours. After replacing the PFA solution with DPBS, the microfluidic chip was disassembled, and cardiac tissue was retrieved and embedded in paraffin. Thin sections of 4 μm thickness were prepared, followed by deparaffinization. The sections were then immersed in a tris-EDTA solution with pH 9.0 for antigen retrieval, achieved by microwave heating at 700W for 20 minutes.

Subsequently, blocking was conducted at room temperature for 1 hour using a 1.5% BSA solution, followed by three DPBS washes. Following this, 100 μl of a staining solution, comprising mouse anti-cTnT primary antibody diluted 1:100 in the blocking buffer, was applied to the tissue section and incubated for 1 hour. After three DPBS washes, 100 μl of a staining solution, containing goat anti-mouse secondary antibody conjugated with Alexa Fluor 488 (1:1000 dilution), ActinRed 555 ReadyProbes Reagent (1:50 dilution), and NucBlue Live ReadyProbes Reagent (1:50 dilution; Cat. #: R37605, Invitrogen, USA), was applied to the tissue section and incubated at room temperature for 1 hour. Subsequent to DPBS washing, the tissue section was mounted with ProLong Gold antifade mountant (Cat. #: P10144, Invitrogen, USA) using a coverslip. Fluorescence images were acquired using the FV1000 confocal microscope.

### Statistical Analysis

Statistical analyses, including sigmoid curve fitting of the dose-response curve for NA and estimation of the EC50 value, were conducted using GraphPad Prism (version 9.5; GraphPad Software, USA). Non-paired Student’s t-tests were employed for the assessment of cell directional analysis, CD31 expression levels, myocardial contractility, and myocardial marker expression levels. For the evaluation of vascular endothelial cell layer permeability and CaT analysis in cardiac tissue, one-way analysis of variance with Tukey’s post-test was applied. Data are presented as means ± standard error of the mean. A p-value less than 0.05 was considered statistically significant.

## Acknowledgment

We express our gratitude to the Central Research Laboratory at Okayama University Medical School for their valuable assistance in preparing paraffin-embedded tissue sections. This research received support from the Japan Society for the Promotion of Science (JSPS) through Grant-in-Aid for Scientific Research (B) (No. 20H04518) and Grant-in-Aid for Scientific Research (A) (No. 21H04960).

